# Elevated FAM84B promotes cell proliferation via interacting with NPM1 in esophageal squamous cell carcinoma

**DOI:** 10.1101/2022.01.10.475754

**Authors:** Fang Wang, Caixia Cheng, Xinhui Wang, Fei Chen, Hongyi Li, Yan Zhou, Yanqiang Wang, Xiaoling Hu, Pengzhou Kong, Ling Zhang, Xiaolong Cheng, Yongping Cui

## Abstract

Family with sequence similarity 84, member B (FAM84B) is a significant copy number amplification gene in the 8q24.21 locus identified by our previous WGS study in esophageal squamous cell carcinoma (ESCC). However, its clinical relevance and potential mechanisms have been elusive. Here, we performed the association analyses between FAM84B_Amp_ and clinicopathological features using our dataset with 507 ESCC samples. The results indicated that, compared with the FAM84B_non-Amp_ patients, the FAM84B_Amp_ patients showed a more aggressive and a worse prognosis. Significant correlation was discovered between the expression level of FAM84B and FAM84B_Amp_ in ESCC cohort. Furthermore, we found that the forced expression change of FAM84B can influence ESCC cell proliferation and cell cycle status, which is probably mediated by NPM1. A direct interaction between FAM84B and the C-terminal (189-294aa) of NPM1 was identified, which increased the NPM1 nuclear expression. Over-expression of NPM1 could inhibit the CDKN2A protein expression, which might affect the ESCC cell cycle. Our results indicate FAM84B CNA may be a potential diagnostic and therapeutic biomarker in ESCC, meanwhile, reveal a novel mechanism of FAM84B that it promotes tumorigenesis via interacting with NPM1 and suppressing CDKN2A.

## Introduction

As one of the two main histological types of esophagus cancer, ESCC shows a higher incidence than EAC in the Chinese population[1]. According to the most recent data, ESCC is the third most prevalent malignant cancer and also the fourth leading cause of cancer death in China[2, 3]. In recent years, progress has been made in improving both diagnostic and therapeutic strategies for ESCC. However, ESCC at advanced stages still has a poor prognosis[4].Therefore, identification of the new therapeutic targets is essential for improving the management of ESCC patients.

In our previous study, we performed 14 WGS and 90 WES using ESCC fresh tumor and matched adjacent normal specimens, respectively. A total of 126 significantly altered regions were identified by FCNAs analysis using GISTIC. It showed 8q24.13-q24.21 was one of the most amplified regions, which contained FAM84B. Amplification of FAM84B was found in 44% and it showed high expression in 57% of the 104 patient samples[5].

FAM84B gene is also known as LRATD2. It is located on chromosome 8q24.21, where the susceptibility locus has been identified in various cancer types[6]. Accumulated evidence has been found to support the association between FAM84B and carcinogenesis. FAM84B is involved in the formation of DNA-repair complexes[7]. It has been identified that FAM84B copy number amplified and promoted tumorigenesis in various cancers. Over-expression FAM84B significantly promoted cell invasion, growth of xenografts and lung metastasis in prostate cancer cells[8]. FAM84B copy number amplification promotes tumorigenesis through the Wnt/β-catenin pathway in pancreatic ductal adenocarcinoma[9]. FAM84B promoted tumor via affecting the Akt/GSK-3β/β-catenin pathway in human glioma[10]. However, its role in ESCC remains unknown.

Here, we analyzed the relationship between FAM84B CNA and clinicopathological characteristics using 507 WGS data of ESCC. The positive correlation between FAM84B CNA and mRNA expression was analyzed in 155 RNA sequencing and TCGA datas. Furthermore, we verified the oncogenic role of FAM84B and elaborated on its potential mechanisms that the complex formation of FAM84B-NPM1 increased the NPM1 nuclear expression which inhibited the CDKN2A protein expression and accelerated cell proliferation via regulating cell cycle in ESCC. Our findings suggested that FAM84B may be not only a novel diagnostic marker and but also a therapeutic target for ESCC.

## Methods

### Samples and clinical data

All the 507 ESCC tumor samples and adjacent normal tissues with good quality and sufficient quantity for in-depth pathological and molecular investigation were obtained from Shanxi and Xinjiang provinces, China. The 507 pairs samples were diagnosed as ESCC by two pathologists independently using hematoxylin and eosin (H&E) staining. Medical records and survival data were obtained for all 507 of ESCC patients. The clinical, epidemiological or pathological features were showed in Table 1. The ESCC individuals were staged according to American Joint Commission for Cancer (AJCC)/International Union Against Cancer (UICC) TNM staging system (the 8th). The study was approved by the Institutional Reviewing Board (IRB) and the Research Committee of Shanxi Medical University.

**TABLE 1.** Association between the copy number amplification of FAM84B levels and clinicopathological variables in ESCC patients.

| Clinic features | FAM84B<sub>Amp</sub><br>≥ 0.5 | FAM84B<sub>non-Amp</sub><br>< 0.5 | *p*-value<sup>a</sup> |
| --- | --- | --- | --- |
| All cases | 109 (21.50%) | 398 (78.50%) |  |
| Age |  |  |  |
| <60 | 51 (23.29%) | 168 (76.71%) | 0.53 |
| ≥60 | 58 (20.14%) | 230 (79.86%) |  |
| Sex |  |  |  |
| female | 38 (20.09%) | 134 (77.91%) | 0.90 |
| male | 71 (21.19%) | 264 (78.81%) |  |
| Location |  |  |  |
| upper | 8 (30.77%) | 18 (69.23%) | 0.41 |
| middle | 65 (20.19%) | 257 (79.81%) |  |
| lower | 36 (22.64%) | 123 (77.36%) |  |
| Smoking status |  |  |  |
| never | 56 (22.13%) | 197 (77.87%) | 0.81 |
| yes | 53 (20.87%) | 201 (79.13%) |  |
| Drinking status |  |  |  |
| never | 76 (21.97%) | 270 (78.03%) | 0.79 |
| yes | 33 (20.5%) | 128 (79.5%) |  |
| Histological grade |  |  |  |
| Grade1 | 8 (17.78%) | 37 (82.22%) | 0.81 |
| Grade2 | 76 (21.78%) | 273 (78.22%) |  |
| Grade3 | 25 (22.12%) | 88 (77.88%) |  |
| Pathologic Stage |  |  |  |
| I&II | 52 (16.25%) | 268 (83.75%) | 0.0003 |
| III | 57 (30.48%) | 130 (69.52%) |  |
| Prognosis (Log-rank test) |  |  |  |
| Deceased | 63 (28.90%) | 155 (71.10%) | 0.0011 |
| Living | 42 (15.38%) | 231 (84.62%) |  |
| Missing | 4 (25%) | 12 (75%) |  |

<sup>a</sup> chi-square test.

### Cell lines and Cell culture

ESCC cell lines used in the research were purchased from Cell Bank of Type Culture Collection of Chinese Academy of Sciences, including KYSE150, KYSE180, KYSE450 and TE-1. ESCC cell lines were cultured in RPMI-1640 medium supplementary (Hyclone) with 10% fetal bovine serum (Gibco), 100 U/ml penicillin, and 100 μg/ml streptomycin. The cell line 293T was from our lab, which was cultured in DEME/HIGH GLUCOSE medium (Hyclone) with 10% fetal bovine serum (Gibco), 100 U/ml penicillin, and 100 μg/ml streptomycin. All ESCC cell lines were cultured at 37°C in a humidified atmosphere with 5% CO_2_. According to the cell state, cell culture medium was replaced. When the cells fusion was about 80-90%, The cells were subcultured.

### MTT assay

5,000 cells per well were plated into 96-well plates and cultured at normal condition for 24 h, 48 h, 72 h and 96 h, respectively. Then 20 μl of 5 mg/ml of MTT (Invitrogen) were added into each well for 4 h at 37°C, until crystals were formed. Then 200 μl DMSO was used to dissolve the crystals and measured the absorbance at 490 nm. The DMSO-treated be seem as control.

### Colony-forming assay

800 cells per well were seeded in 6-well plates and cultured conventionally for 10 days. In the end, the colonies were fixed in 4% paraformaldehyde for 30 min and stained with 1% crystal violet for 20 min at room temperature. The colonies containing more than 50 cells were photographed and counted.

### Flow cytometry

For cell-cycle profile analysis, transfected cells were digested with trypsin into single-cell suspensions, and 1×10^6^ suspended cells were collected for experiments. The collected cells were washed with PBS three times, following by incubating the cells for at least 15 min with 1 ml of propi-dium iodide (PI) dyeing liquid (KeyGen Biotech Co., China), and analyzed by flow cytometry (BD Biosciences, USA).

### Plasmids construction and transfections

FAM84B-FL construct was made using pGEX-5X-1 vector with the BamH1 and EcoR1 sites. The NPM1 deletion-mutants (NPM1-FL, NPM1-1-117aa, NPM1-118-188aa, NPM1-189-294aa, NPM1-1-188aa, NPM1-118-294aa) with a HA tag and the plasmid of pCMV-FLAG-FAM84B with a FLAG tag were purchased from PolePolar Biotechnology Co (Beijing, China). The siRNA (RiboBio) of NPM1 was used to knockdown NPM1. The siRNA sequences are: si-NPM1-RNA1: 5’-ACTGCTTTATACTTTGTCA-3’; si-NPM1-RNA2:AATGGCAAATAGTCTTGTA-3’; The lentivirus for stable over-expression and knock down of FAM84B gene were constructed and packaged by Hanbio Biotechnology Co. (Shanghai, China). For knockdown of endogenous FAM84B, we used vectors containing the sequence: FAM84B-shRNA1: 5’-CACCTAAGTTACAAGGAAGTTCTCGAGAACTTCCTTGTAACTTAGGTG-3’; FAM84B-shRNA2: 5’-AGTCTAGAGGACCTGATCATGCTCGAGCATGATCAGGTCCTCTAGACT-3’. Plasmids were performed via the lipofectamine™ 2000 transfection reagent (Invitrogen) according to the manufacturer. The siRNA was transfected with riboFect™ CP Transfection Kit (C10511-1).

### Real-time quantitative PCR (qPCR)

qPCR was used for measuring mRNA expression. Total RNA was isolated from cells using the RNA extraction reagent (Takara). Reverse transcription was performed using PrimeScriptTM RT reagent kit (Takara), qPCR was performed using the SYBR Green Premix Ex TaqTM (Takara) and specific primers. All qPCR reactions were performed in triplicate with an Applied Biosystems StepOnePlus (ABI). The relative expression of genes was determined by normalization to GAPDH expression according to the manufacturer’s instructions. Data analysis was performed using the formula: 2^-ΔΔCt^. The primers are listed in Suppl Table 1.

### Western blot

Protein levels of the genes were detected through western blot. Briefly, Cells were lysed using RIPA buffer for 1 h on ice, the protein concentrations were determined via a BCA assay kit (Boster, Wuhan). The proteins were separated by SDS-PAGE. 50 μg of protein and 4×loading buffer were boiled for 10 min and separated by SDS-PAGE (10% separating gel and 5% stacking gel), the proteins were transferred onto polyvinylidene fluoride (PVDF) membranes (Millipore, USA) that were subjected to blocking by 5% skimmed milk for 1 h at room temperature, then incubated with the specific antibodies at 4°C overnight. The goat anti-mouse and goat anti-rabbit second antibodies were got from Odyssey. Relative amount of gene product was normalized to GAPDH levels.

Proteins were detected by using anti-FAM84B (Proteintech, 18421-1-AP), anti-NPM1 (Proteintech, 60096-1-Ig), anti-CCND1 (Proteintech, 60186-1-Ig), anti-CDK4 (Proteintech, 11026-1-AP), CDK6 (Proteintech, 14052-1-AP), anti-FLAG (Cell Signaling, #14793), anti-HA (Abcam, 9110), anti-GST (Proteintech, 66001-2-Ig), anti-pRb (Cell Signaling, #8516), anti-CDKN2A (10883-1-AP), anti-E2F1 (12171-1-AP) and anti-GAPDH (Proteintech, 60004-1-Ig).

### Immunofluorescence

Adherent cells were seeded in six-well plates with chamber slides 1day before immunofluorescence. After this, the cells were fixed and permeabilized at room temperature. Then, the cells were incubated with primary anti-FAM84B (1:50, Proteintech) and anti-NPM1 (1:50, Proteintech) at 4°C overnight. After washing with PBST, cells were incubated with Alexa FluorTM 594 anti-rabbit antibody and Alexa Fluor® 488 anti-mouse antibody (Invitrogen) of 2 drops/ml for 30min at room temperature. Coverslips were mounted on slides with ProLong(tm) Diamond Antifade Mountant with DAPI (Thermo Fisher) according to the manufacturer’s instructions. Fluorescent images were taken using an LSM700 confocal laser scanning microscope (200×).

### Mass spectrometry analysis and CO-IP

The IP assay was essentially done as described. Briefly, KYSE450 cells were infected with the plasmid of pCMV-FLAG-FAM84B. 1□mg of protein with anti-FAM84B or anti-IgG (Proteintech, 10284-1-AP) and Protein A/G plus-agarose (Santa Cruz Biotechnology) overnight at 4°C. Meanwhile, suitable proportion cell lysates were stored as input. Then washing three times, the antibody/lysate mixture was captured by the Protein A/G plus-agarose and detected by electrophoresis. Finally, we found out the differential expression bands for Mass spectrometry analysis and sequence (Shanghai Applied Protein Technology co. ltd).

For Co-IP Assay, the cell lysates containing 1□mg of protein with individual antibodies and Protein A/G plus-agarose (Santa Cruz Biotechnology) overnight at 4°C. Beads were washed and eluted with sample buffer, and boiled for 10 min at 100°C and centrifuged for western blot. The supernatants were run on SDS-PAGE and blotted with respective antibodies. Antibodies used for IP were anti-FLAG (Cell Signaling, #14793) and anti-FAM84B (Proteintech, 18421-1-AP). Antibodies used for the western blot were anti-HA (Abcam, 9110) and anti-NPM1 (Proteintech, 60096-1-Ig). Anti-IgG was used as a negative control.

### GST-pull down assay

The GST pull-down assay was essentially done as described by Sun et al. The fusion proteins with GST-tag were expressed in Bl-21 following induction with IPTG. Purified 50µg fusion proteins were immobilized on 20 µl GSH-sepharose for 2 hours at 4 °C, then washed with PBS-T binding buffer (PBS, pH 7.4, 1% Tween 20) three times. Immobilized proteins were incubated for overnight at 4 °C with 5 µg of NPM1 product from an *in vitro* translation reaction. The plasmid of pCMV-NPM1-HA was transfected via the lipofectamineTM 2000 transfection reagent according to the manufacturer. After this, beads were washed four times with lysis buffer and 4× loading buffer was added to beads and boiled for 10 minutes and centrifuged for western blot. The supernatants were run on SDS-PAGE and blotted with respective antibodies. Antibodies used for GST pull-down assay were anti-GST, anti-HA anti-GAPDH.

### Mouse xenograft assay

The mouse xenograft assay was performed as described previously. Briefly, 3 × 10^6^ KYSE150 cells with stably knock-down of FAM84B and control vector were suspended in PBS and injected subcutaneously into 6-week-old BALB/c nude mice (Beijing, China). The tumor size was measured every four days and calculated. After 4 weeks, tumors were removed and weighed, snap frozen in liquid nitrogen. Tumor size was measured and presented as mean±Standard Deviation (SD). Tumor volume calculations were obtained using the formula V= (W^2^×L)/2. For animal studies, approval was obtained from the appropriate animal care committee of Shanxi Medical University.

### Immunohistochemistry (IHC)

The formalin-fixed paraffin-embedded xenograft tumor tissues were immunohistochemically stained. Next antigen retrieval and non-specific antigen blocking, section were incubated with the first antibody for overnight at 4°C. Added the second antibody and incubated 30-40 min. DAB plus kit (MaiXin, Fuzhou,China) was used to develop the staining. The nuclear amount of proteins was analyzed with Aperio Nuclear v.9 software. Statistical analyses were performed with GraphPad Prism 7.0. Proteins were detected by using anti-Ki-67 (Proteintech), anti-FAM84B (Proteintech, 18421-1-AP), anti-NPM1 (Proteintech, 60096-1-Ig), anti-CCND1 (Proteintech, 60186-1-Ig), anti-CDK4 (Proteintech, 11026-1-AP).

### Focal copy number alterations analysis

Sample selection was based on CNAs by GISTIC2.0 (log2 ratio ≤ 0.5 for deletions and >0.5 for gains). The FAM84B copy number alterations in 32 types cancers were downloaded via cBioPortal for Cancer Genomics in the TCGA database (https://www.cbioportal.org). The correlation analysis between FAM84B copy number amplification and mRNA expression was downloaded from TCGA via xenabrowser (https://xenabrowser.net/heatmap/). The 507 pairs ESCC tissue were carried out WGS sequencing. All FASTQ files are going to be uploaded to Genome Sequence Archive (GSA) in Beijing Institute of Genomics (BIG) Data Center, the accession number is HRA000021, that will be publicly accessible at http://bigd.big.ac.cn/gsa.

### Statistical analysis

All experiments were done in triplicates and data were presented as mean ± SD or ± SEM. Data from two groups were analyzed by unpaired t-test and more than two groups were analyzed by one-way ANOVA. The correlation between FAM84B copy number amplification and the clinical variables in ESCC was analyzed by Chi-square test. Kaplan–Meier analysis and Log rank test or Breslow test were used for survival analysis. Cox proportional hazards regression model was used for multivariate survival analysis. Statistical Package for Social Science for Windows (SPSS20.0, USA) was used for all statistical analysis. The correlations between FAM84B copy number amplification and FAM84B gene expression were performed using nonparametric correlation (Spearman) by GraphPad prism software. *P* < 0.05 was considered to be statistically significant.

## Results

### FAM84B copy number amplification is correlated with prognosis in ESCC

We focused on somatic FCNAs characterized by GISTIC2.0 in the 507 ESCC patients. It showed high-amplitude copy number changes in 8q24.21 which including FAM84B. FAM84B was defined as an amplified gene in 109 out of 507 tumors (21.5%, Fig. 1A and 1B). Furtherly, we analysed the relationship between FAM84B copy number amplification and clinicopathological characteristics in ESCC. The 507 ESCC patients was divided into two groups: patients with copy number amplification of FAM84B (named as FAM84B_Amp_, ≥ 0.5) and patients without copy number amplification of FAM84B (named as FAM84B_non-Amp_,< 0.5). The results showed FAM84B_Amp_ was associated with the invasion depth (T stage) (*P* < 0.001) and survival status (*P* = 0.0011) in the ESCC patients (Table 1). Kaplan–Meier survival analysis showed the patients with FAM84B_Amp_ had a shorter survival time than those with FAM84B_non-Amp_ (*P* < 0.001, Fig. 1C). The cox multivariate analysis showed that T stage (HR = 2.301, 95 % CI: 1.744-3.037, *P* < 0.001), Grade (HR = 0.57, 95 % CI: 0.363-0.896, *P* = 0.015) and FAM84B_Amp_ (HR = 0.649, 95 % CI: 0.482-0.874, *P* = 0.004) were predictive factors for overall survival, respectively (Fig. 1D). Moreover, the ESCC patients can be divided into four groups with different survival rates according to the FAM84B_Amp_ and T stage status (Fig. 1E and 1F, Suppl. Table. 2). These results suggested that the patients with FAM84B_Amp_ had a deeper invasion and a worse prognosis. FAM84B_Amp_ might play an important role in the tumorigenesis and development of ESCC. Meanwhile, FAM84B_Amp_ was correlated with the survival status in the patients with female (*P* = 0.007), male (*P* =00.017), age <60 (*P* = 0.002), no smoking (*P* = 0.001), no drinking (*P* < 0.001), T stage [(*P* = 0.001) and grade=3 (*P* = 0.001), respectively (Suppl. Fig. 1 and 2).

**Fig 1.**
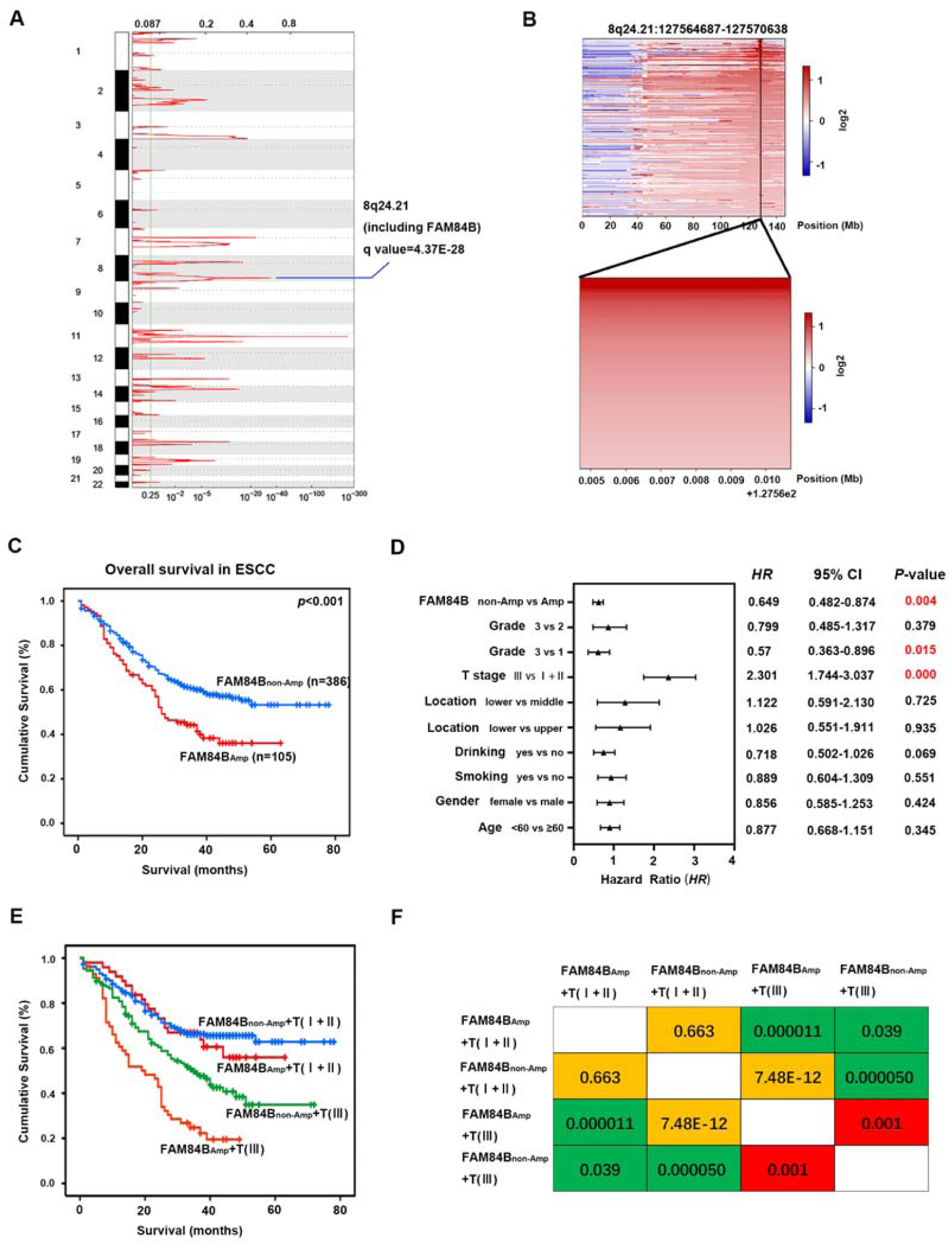
FAM84B_Amp_ is correlated with the prognosis in 507 ESCC patients. (A) The significant focal SCNA characterized by GISTIC in the 507 ESCC cohort. (B) Heatmap of CNA log2 ratio of read coverage through 109 ESCC individuals in 8q24.21 and FAM84B regions (upper) and detected significant amplification of FAM84B (bottom). (C) Kaplan-Meier survival plot showed the patients with FAM84B_Amp_ had worse survival than those with FAM84B_non-Amp_ (*P* < 0.001). (D) Multivariate analysis by cox proportional hazards regression model for overall survival in 507 ESCC patients. (E) Combination of FAM84B_Amp_ and T stage can effectively divide the 507 ESCC patients into four groups which have different survival rates. (F) Pairwise comparison matrix of the four groups divided by combination of FAM84B_Amp_ and T stage, the Log Rank *P* values were shown.

To investigate FAM84B CNAs in various cancer types, we examined the patterns of FAM84B amplification in 10,802 tumor samples belonging to a total of 32 cancer types (TCGA dataset). As shown in Fig. 2A, 28 tumor types displayed amplification to various degrees of FAM84B. Consistent with our results, the pan-cancer patients with FAM84B_Amp_ had significantly shorter overall and relapse-free survival compared with the wild type patients (*P* = 0.0233; *P* = 7.52e-9, Fig. 2B), suggesting that FAM84B amplification might result in a worse prognosis. Additionally, we conducted the correlation analysis between the FAM84B_Amp_ and the RNA expression level in ESCC databases. Consistent with paired ESCC samples, the positive correlation was found in both TCGA ESCC cohort (r = 0.449; *P* = 0.0011; n = 95; Fig. 2C) and 155 RNA-seq ESCC cohort (r = 0.449, *P* < 0.001; n = 155; Fig. 2D). Meanwhile, immunohistochemistry analysis of FAM84B in 104 primary ESCC samples showed that the expression of FAM84B was markedly higher in tumors than the matched normal tissues[11]. These results speculated that FAM84B_Amp_ and high-expression might participate in the progress of ESCC and FAM84B might serve as a biomarker for prognosis of ESCC patients.

**Fig 2.**
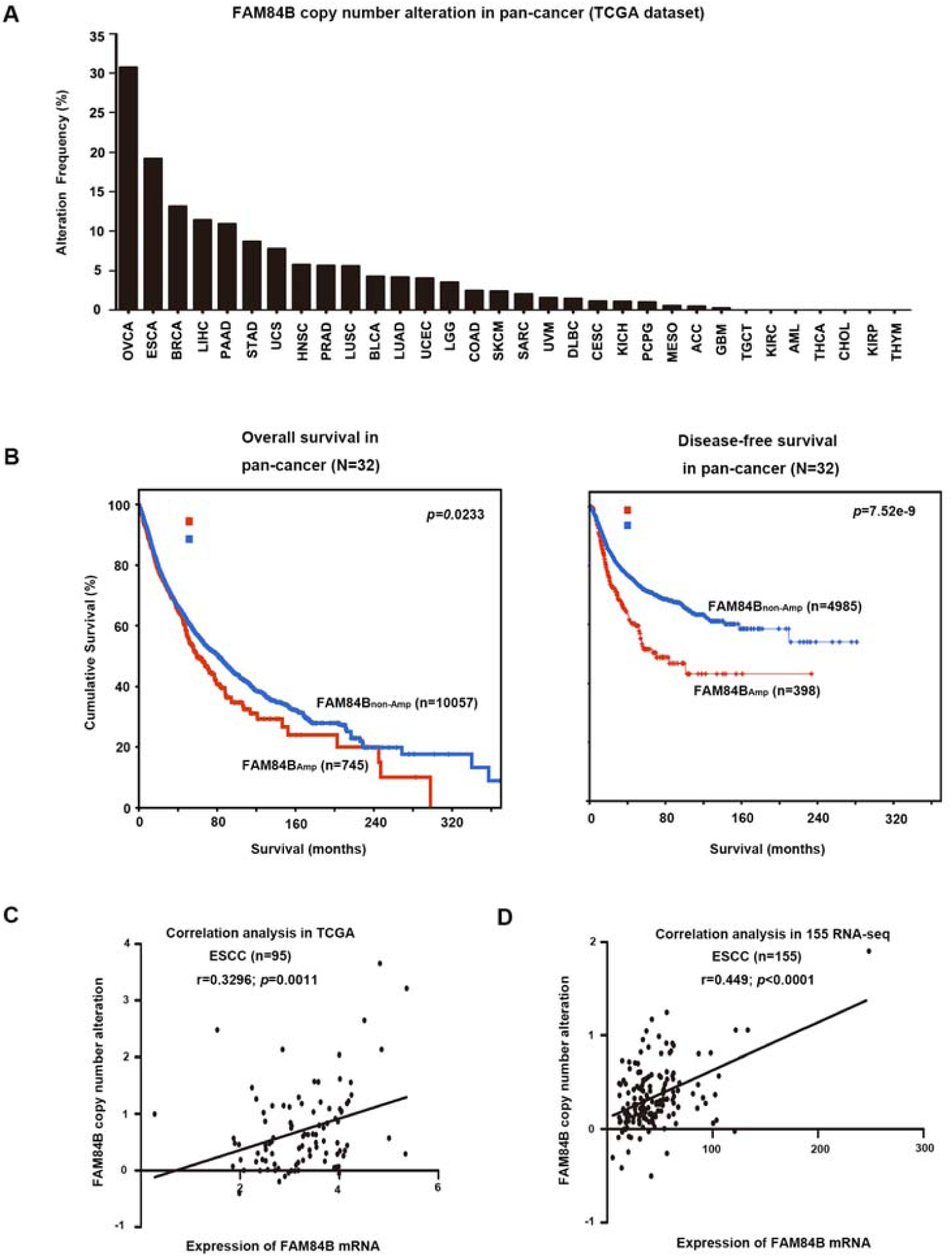
Copy number amplification and expression of FAM84B in pan-cancer. (A) GISTIC2.0 analysis 32 human cancer types (The Cancer Genome Atlas, TCGA) showing varying degrees of copy number amplification of FAM84B. (B) FAM84B_Amp_ was correlated with overall and relapse-free survival in the TCGA pan-cancer cohort (*P* = 0.0233; n = 10802; *P* = 7.52e-9; n = 5383; N = 32 cancer types). (C-D) The correlation analysis of FAM84B_Amp_ and expression in TCGA (left, *P* = 0.0011; n = 95) and 155 ESCC cohort (right, *P* < 0.001; n = 155).

### FAM84B promoted ESCC proliferation and cell cycle *in vivo* and *in vitro*

To elucidate the biological effect of FAM84B copy number amplification in ESCC, the protein levels of FAM84B in ESCC cell lines were tested, including the immortal embryonic esophageal epithelium cell lines NE-2, and ESCC cell lines KYSE180, KYSE150, KYSE450 by quantitative real-time PCR (Suppl. Fig. 3a). Of these cell lines, KYSE450 cell line was selected for over-expression experiments, meanwhile, KYSE150 and KYSE180 cell lines were selected for knockdown experiments. The transfection efficiency was detected by western blot assay respectively (Suppl. Fig. 3b and 3c). FAM84B exogenous over-expression significantly increased cell proliferation (Fig. 3A) and colony formation (Fig. 3C) in the KYSE450 cell line. The flow cytometry assay showed over-expression of FAM84B promoted cell cycle (Fig. 3E). Knockdown of FAM84B significantly inhibited cell proliferation (Fig. 3B) and colony formation (Fig. 3D) compared with the control group KYSE150 and KYSE180 cell lines. The flow cytometry study indicated knockdown of FAM84B decreased cell cycle and mainly arrested in the G1/S phase (Fig. 3F).

**Fig 3.**
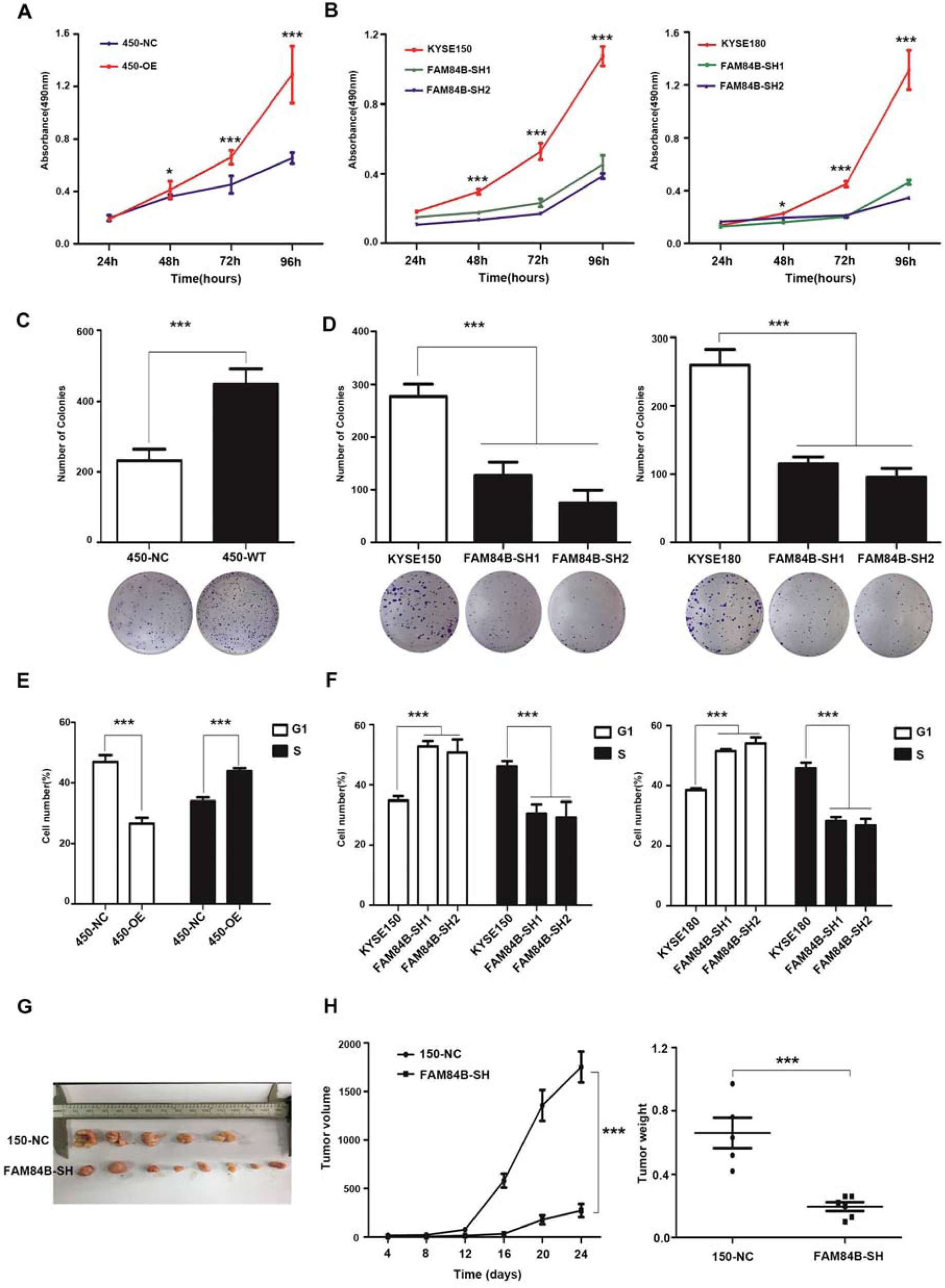
The expression of FAM84B significantly effected ESCC cell proliferation and cell cycle. (A) Over-expression of FAM84B significantly increased the ability of proliferation in KYSE450. (B) Knock-down FAM84B dramatically decreased the ability of proliferation in KYSE150 and KYSE180 cells. (C) Over-expression of FAM84B significantly promoted the the ability of colony formation in KYSE450. (D) Knock-down FAM84B inhibited the ability of colony formation in KYSE150 and KYSE180 cells. (E) Over-expression FAM84B promoted cell cycle by flow cytometry in KYSE450 cells. (F) Knock-down FAM84B inhibited cell cycle and arrests to the G1/S phase in KYSE150 and KYSE180 cells. (G-H) FAM84B knock down markedly inhibited tumor growth and decreases the weight of the tumor mass in the xenograft system. Statistical analysis is performed with one-way ANOVA. \**P* < 0.05, \*\*\**P* < 0.001.

To further confirm this conclusion *in vivo*, we established a subcutaneous transplantation tumor model in female BALB/c nude mice using the KYSE150 cells with stably knock-down of FAM84B and control vector. Four weeks later, tumors were removed and weighed. The results shown that the tumor volume of FAM84B knock-down group was smaller than the control group (t - test, *P* < 0.001). The tumor weight of FAM84B knock-down group was lighter than the control group (t - test, *P* < 0.001) (Fig. 3G and Fig. 3H). These results suggested that FAM84B might as an oncogene promoted ESCC tumorigenesis via regulating cell cycle *in vitro* and *in vivo*.

### NPM1 may be a candidate target gene of FAM84B in ESCC

The significant cell phenotypes changes lead us to investigate the interrelation of FAM84B with tumorigenesis in ESCC. IP/MS experiment was performed to explore the FAM84B-associated protein(s). Briefly, we transfected pCMV-FLAG-FAM84B plasmid in KYSE150 cell line. The cell lysate was subjected to CO-IP assay, and bound proteins were subjected to silver staining (Suppl. Fig. 4a). NPM1 is identified as an interacting protein of FAM84B by MS (Suppl. Fig. 4b). NPM1, also known as B23 protein, a multifunctional phosphoprotein, resided primarily in the the granular regions of the nucleolus. NPM1 protein can shuttle between the nucleus, the nucleoplasm, and the cytoplasm during the cell cycle[12]. NPM1 contained a number of motifs that mediated the interactions with the binding partners and affected its cellular localization[13]. To determine whether FAM84B and NPM1 co-localize with each other, immunofluorescence analysis targeting FAM84B and NPM1 was performed in KYSE150 cells. The result showed that FAM84B and NPM1 are associated with each other (Fig. 4A). Furthermore, CO-IP assays were conducted to examine the endogenous and exogenous interaction of FAM84B and NPM1 (Fig. 4B and 4C). Meanwhile, the GST-pull down assay results showed FAM84B can bind to NPM1 and form a complex structure in *vivo* (Fig. 4D). These results suggested that NPM1 is specifically co-immunoprecipitation with FAM84B, and may be a candidate target gene of FAM84B in ESCC.

**Fig 4.**
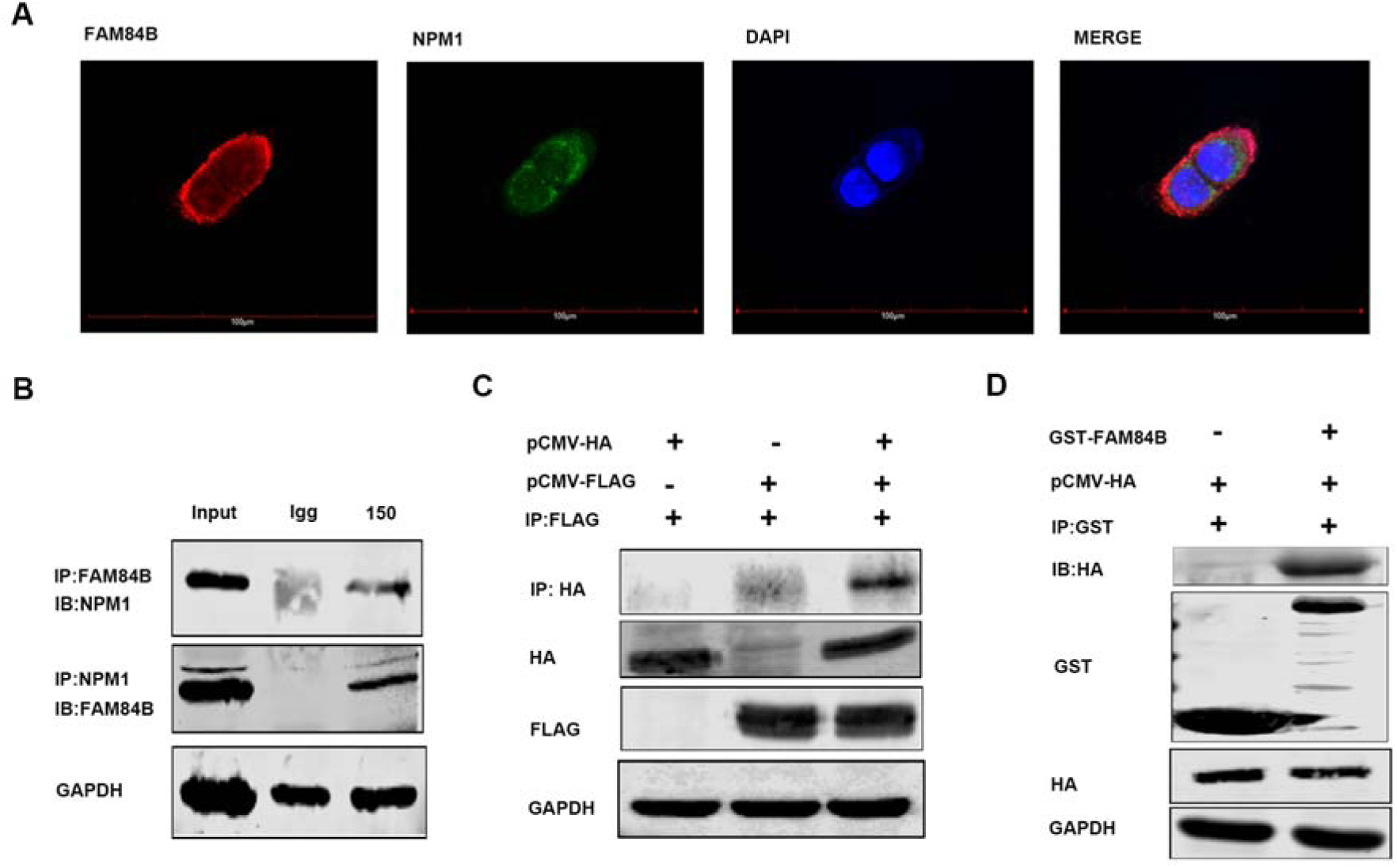
FAM84B interacted with NPM1 in ESCC cells. (A) The co-location of endogenous FAM84B and NPM1 by immunofluorescence in KYSE150 cells. The first image of cells was stained with the first antibody of anti-FAM84B and the second antibody of Alexa FluorTM 594 goat anti-rabbit (red); the second image of cells were stained with the first antibody of anti-NPM1 and Alexa Fluor® 488 donkey anti-mouse (green); the third image nuclei of cells were stained with DAPI (blue); the last image was merged (yellow). (B) The interaction with FAM84B and NPM1 were detected in endogenous cells by CO-IP assay. (C) The interaction with FAM84B and NPM1 were detected in exogenous cells by CO-IP assay. (D) The interaction directly with FAM84B and NPM1 were detected in 293T cells by GST-pull down assay.

### FAM84B regulates NPM1 expression through binding to the target regions of NPM1

In previous studies, NPM1 contained three functional domains: an N-terminal oligomerization domain (OligoD) bearing chaperone activity, the C-terminal nucleic acid binding domain (NBD), and two central acid domains for histone binding (HistonD)[14]. To further explored the interaction structure domain of NPM1 and FAM84B, we mapped the domains of NPM1 using a series of HA-tagged NPM1 deletion-mutants (1-117aa, 118-188aa, 189-294aa, 1-188aa, 118-294aa and full length) fused to HA tag[15]. CO-IP assays revealed that the FLAG-FAM84B bound to NPM1-118-294aa and the full-length NPM1, but not to NPM1-1-117aa, NPM1-118-188aa, and NPM1-1-188aa. The results indicated that the C-terminal (189-294aa) is the target regions of NPM1 for its interaction with FAM84B (Fig. 5A and 5B). Consistently, ectopic expression of FAM84B increased the level of NPM1 in a dose-dependent manner (Fig. 5C). Meanwhile, we detected the NPM1 expression change in FAM84B over-expression cells. Nucleoplasmic separation assay showed FAM84B over-expression increased the nuclear localization and expression of FAM84B and NPM1 (Fig. 5D). These results indicated FAM84B might interact with NPM1 through targeting the NPM1-118-294aa domain, which increased the expression of NPM1 in the cell nuclear.

**Fig 5.**
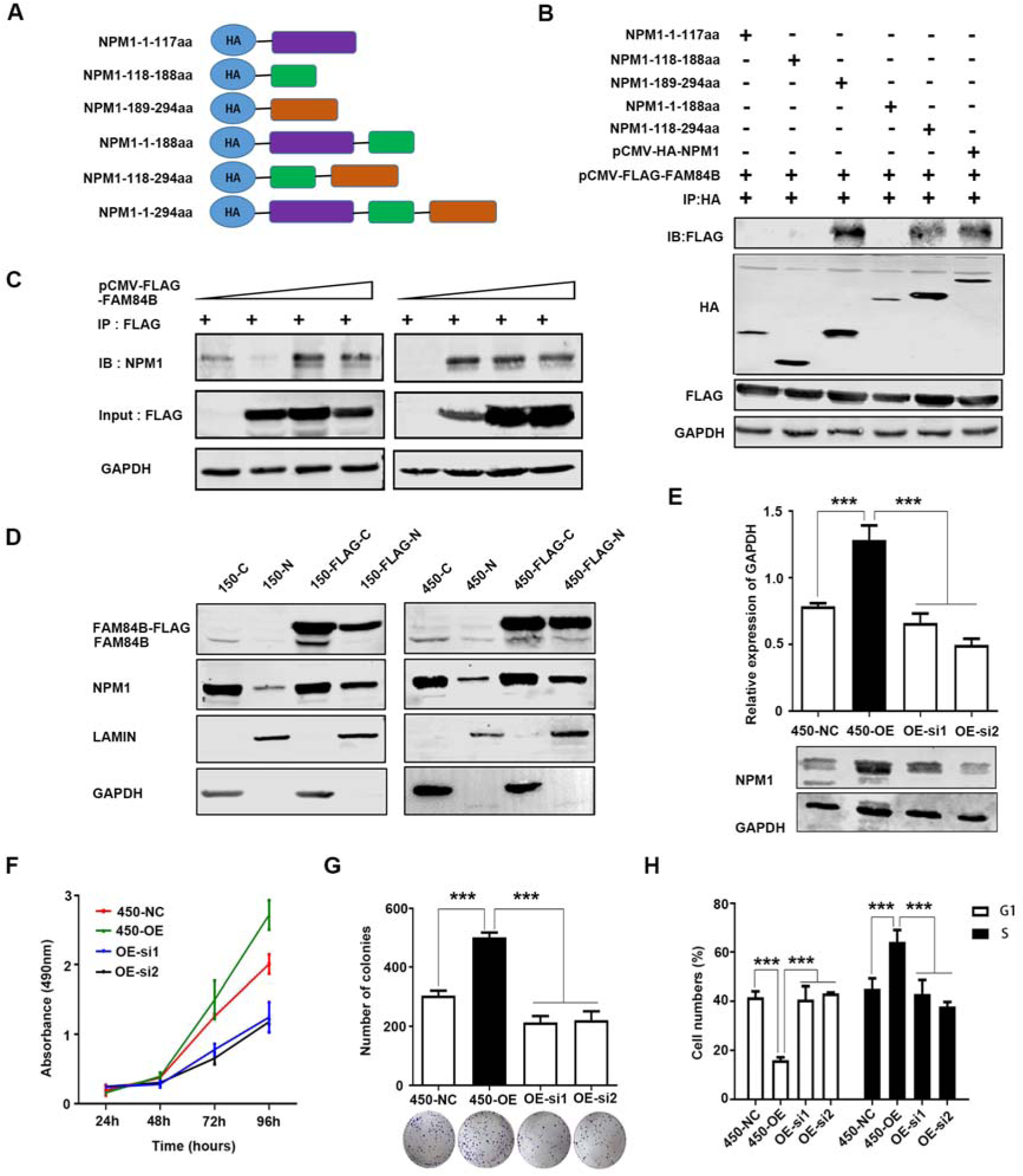
NPM1 may be a downstream target of FAM84B in ESCC. (A) Schematic representation of full-length and deletion mutants of HA-tagged NPM1 protein. (B) The FLAG-FAM84B protein were incubated with full-length and deletion mutants of HA-tagged NPM1 protein using CO-IP assay. (C) The NPM1 was examined after transfecting pCMV-FLAG-FAM84B in gradient amount. (D) The NPM1 nuclear expression was detected in FAM84B over expression cells. (E) Detected the efficiency of knock-down NPM1 in stably over-expression FAM84B cells by western blot. (F) Knock-down NPM1 inhibited the capability of cell proliferation in stably over-expression FAM84B cells. (G) Knock-down NPM1 inhibited the capability of colony formation in stably over-expression FAM84B cells. (H) Knock-down NPM1 inhibited cell cycle in stably over-expression FAM84B cells. Statistical analysis was performed using one-way ANOVA. \*\*\**P* < 0.001.

### NPM1 may be a downstream target of FAM84B in ESCC

NPM1 is an abundant nucleolar protein that is involved in not only a variety of biological processes but also the pathogenesis of several human malignancies[16, 17]. To investigate its function in ESCC, knockdown and over-expression NPM1 experiments were performed in ESCC cell lines respectively. The transfection efficiency was detected by western blot assay (Suppl. Fig. 5a and 5b). Interestingly, the result showed knock-down of NPM1 significantly decreased cell proliferation, colony formation ability and cell cycle. Meanwhile, over-expression of NPM1 markedly increased cell proliferation, colony formation ability and cell cycle. These result indicated that NPM1 maybe as a oncogene promoted tumor formation in ESCC (Suppl. Fig. 5c, 5d, 5e).

To confirm whether FAM84B promoted cell cycle through NPM1, we carried out the interference and rescue experiment of NPM1. Knockdown NPM1 was used to detected a series of phenotype changes in FAM84B over-expression ESCC cells (Fig. 5E). The results showed that the forced knock-down of NPM1 inhibited the ability of cell proliferation and colony formation (Fig. 5F and 5G). We also found the forced knock-down of NPM1 increased the proportion of G1phase cells and decreased the proportion of S phase cells (Fig. 5H). These results indicated that NPM1 inhibition can reverse a series of phenotype changes caused by FAM84B high-expression. It was confirmed further that NPM1 may be as a candidate target gene of FAM84B. FAM84B promoted ESCC tumorigenesis by targeting NPM1.

### FAM84B-NPM1 might regulate cell cycle via suppressing the expression of CDKN2A

Although there was a putative FAM84B regulated cell proliferation and tumor growth through NPM1, but the underlying molecular mechanism contributing to cell cycle pathway in ESCC remains unknown. Hence, we screened the MS results of FAM84B and the proteins interacting with NPM1 using NCBI database. As a result, CDKN2A was identified as targeted protein of the FAM84B and NPM1 complex (Suppl. Fig. 6). CDKN2A, a tumor suppressor gene, is located on chromosome 9p21 and has three exons. It events phosphorylation of Rb protein and halts the cell cycle progressing from G1 to S phase[18]. CO-IP assays were performed to confirm the interaction of CDKN2A with FAM84B and NPM1 in cells. The result showed CDKN2A not only interacted with FAM84B, but also interacted with NPM1 (Fig. 6A and 6B). Meanwhile, We detected the expression of CDKN2A after NPM1 over-expression. The result showed that NPM1 over-expression could decrease the CDKN2A expression in 150 and 450 cells (Fig. 6C). Furtherly, we detected the expression of cell cycle protein after FAM84B over-expression/knock-down in cells. The results showed FAM84B over-expression resulted in increased expression of CDK4, CDK6, CCND1, p-Rb and E2F by RT-PCR and westen blot, FAM84B knock-down resulted in a significant decrease of the cell cycle proteins expression in mRNA and protein levels (Fig. 6D and Suppl. Fig. 7). The pattern of cell cycle proteins and mRNA expression in cells promoted us to further investigate the expression of mouse tumors. Therefore, we detected the cell cycle proteins using the nude mouse tumors by IHC assay. Compared with the control group, Ki-67 positive cells were reduced significantly in the FAM84B knockdown group. Knock down FAM84B inhibited the expression of cell cycle proteins in the tumor tissues (Fig. 6E). Similar results of CDK4 and CCND1 expression were observed in *vivo* and in *vitro* studies, supporting that FAM84B-NPM1 might regulate cell cycle pathway via suppressing the expression of CDKN2A.

**Fig 6.**
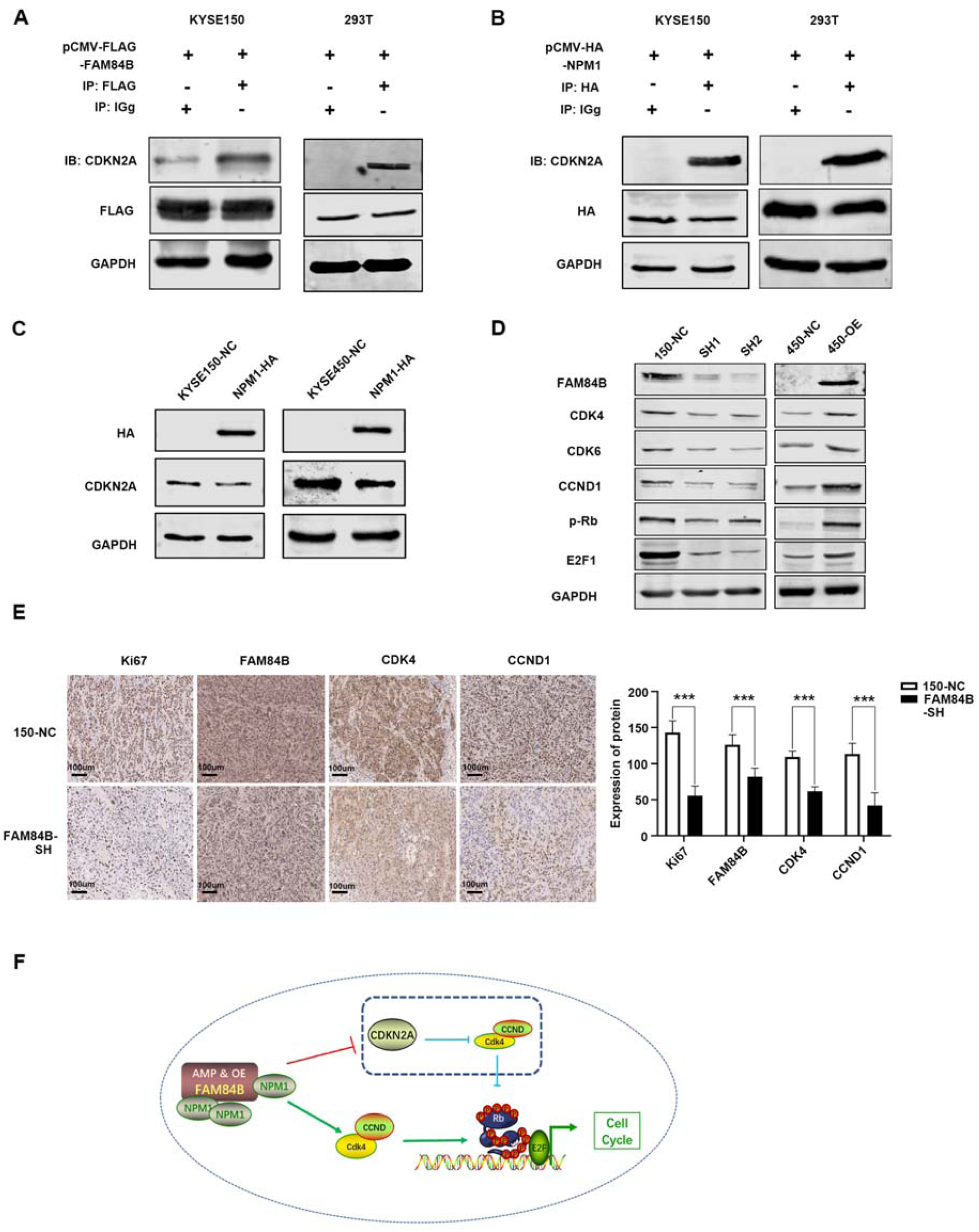
FAM84B-NPM1 might regulate cell cycle via suppressing the expression of CDKN2A. (A) The interaction with FAM84B and CDKN2A were detected in cells by CO-IP assays. (B) The interaction with NPM1 and CDKN2A were detected in cells by CO-IP assays. (C) NPM1 over expression decreased the CDKN2A expression in 150 and 450 cells. (D) Deteced the expression of cell cycle proteins by western blot in over-expression/knock down FAM84B cells. (E) Immunohistochemical images showed the expression level of Ki-67, FAM84B, NPM1, CDK4 and CCND1 from mice injected with KYSE150 NC cells and FAM84B knockdown cells, Magnification, 200×. (F) Diagram showing how copy number amplification of FAM84B contributes to tumorigenesis of ESCC via regulation of cell cycle. Statistical analysis was performed using one-way ANOVA. \*\*\**P* < 0.001,

This study indicated that FAM84B copy number amplification resulted in increased the expression of FAM84B, which was correlated with prognosis in ESCC. Furthermore, FAM84B gene act as an oncogene to promote ESCC tumorigenesis. FAM84B copy number amplification and high expression may accelerate the cell cycle process and promote cell proliferation by binding with NPM1 functional domain in ESCC. FAM84B interacted with NPM1 and increased the expression of NPM1 in the cell nuclear. NPM1 over expression could inhibit the CDKN2A expression. Therefore, the cell cycle process and the cell proliferation of ESCC are inhibited as the expression of CDKN2A is depressed (Fig. 6F). In a word, FAM84B copy number amplification and high expression may be a potential diagnostic and therapeutic biomarker in ESCC.

## Discussion

Currently, the treatment of ESCC relies on surgery, chemotherapy, radiotherapy, or combinations of these, but limited on effective molecularly targeted therapies that may attribute to the precise molecular events underlining ESCC formation remain only partially understood[19, 20]. In this study, FAM84B was frequently amplified in ESCC and FAM84B as oncogenic drivers for ESCC progression. Our findings are consistent with the recently described association about FAM84B in gastroesophageal junction adenocarcinomas, pancreatic ductal adenocarcinoma and prostate cancer[20]. Meanwhile, FAM84B amplification was closely related to invasion depth and worse survival of 507 ESCC patients. A positive correlation between FAM84B CNAs and RNA expression was found in two ESCC cohorts. This finding suggests that FAM84B amplification and the resultant increased levels of FAM84B expression are associated with progression. It indicated that FAM84B_Amp_ and over-expression might a promising marker and target for cancer diagnosis and therapy.

As multiple genetic lesions in oncogenes or tumor suppressors are involved in cancer initiation and maintenance, targeting of these oncogenic pathways could be a very powerful strategy to inhibit tumor growth[21, 22]. We confirmed that the forced expression change of FAM84B can influence ESCC cell proliferation and cell cycle status. FAM84B can interact with NPM1 to form a complex and regulated cell cycle via suppressing the expression of CDKN2A. The FAM84B-NPM1 complex combined CDKN2A and inhibited the expression, which accelerated CCND-CDK4/6 mediated pRb that lead to the release of E2Fs and promoted cell cycle in ESCC tumorigenesis and progression. The CDK4/6-inhibitor of CDK4 (INK4)-retinoblastoma (Rb) pathway plays a crucial role in cell cycle progression and its dysregulation is an important contributor to endocrine therapy resistance[23]. Palbociclib induced cell cycle arrest in G1 phase and decreased cell migration and invasion via CDK4/Rb signaling pathway[24].

Our study has a few limitations. Firstly, CDKN2A/p16 is a tumor suppressor gene locus that lies adjacent to the 9p21.3 genomic region, which is the site of loss of heterozygosity in some malignant tumors[25]. It encoded the tumor suppressor protein p16, which inhibited CCND-CDK4/6 mediated phosphorylation of the Rb protein that, in turn, leads to the release of E2Fs[26]. Studies have found that the copy number of CDKN2A/p16 was deleted in many cancers, including oral squamous cell carcinoma (OSCC)[27], head-neck squamous cell carcinoma (HSCC)[28] and ESCC[29]. However, the copy number of FAM84B was increased in ESCC and others. We identified nearly statistically significant mutual exclusivity between mutations in FAM84B and CDKN2A in various cancer types (Suppl. Fig. 8, *P* = 0.08). Secondly, it will be required clinical validation that the hypotheses generated from this study. To evaluate the candidate predictive biomarkers which identified from this study would be crucial in clinical trials. In summary, we found that a significant portion of ESCC patients had FAM84B copy number amplification and may potentially benefit from targeted therapies. We provided its potential to impact clinical outcomes and therapeutic targets for ESCC treatment. But further efforts were required to exploit this information to develop a prognostic method and to identify therapeutic targets that could be used to treat biomarker-selected groups of patients with ESCC.

## Ethics approval and consent to participate

This study was approved by the Institutional Reviewing Board and the Research Committee of Shanxi Medical University, and written consent was obtained from all participants.

## Consent for publication

No consent was involved in this publication.

## Availability of supporting data

All data that support the findings of this study are available from the corresponding authors upon reasonable request.

## Authors’ Contributions

Yongping Cui and Xiaolong Cheng contributed to conception and design of the study. Fang Wang, Caixia Cheng, Xinhui Wang, Fei Chen organized the database. Fang Wang and Yan Zhou performed the statistical analysis. Fang Wang wrote the first draft of the manuscript. Yanqiang Wang, Hongyi Li, Xiaoling Hu, Pengzhou Kongand Ling Zhang edited the manuscript. Yongping Cui reviewed the manuscript. All authors contributed to manuscript revision, read and approved the submitted version.

## Acknowledgements

This work uses ESCC samples that have been provided the department of Pathology, Shanxi Province Cancer Hospital.

